# Ordinal position in auditory sequences is encoded in population dynamics in macaque frontal cortex

**DOI:** 10.64898/2026.10.08.757540

**Authors:** Steven P. Errington, Beshoy Agayby, Joel I. Berger, Timothy D. Griffiths, Christopher I. Petkov, Yukiko Kikuchi

## Abstract

The ability to extract structure from sensory sequences requires tracking not only stimulus identity but also ordinal position within a sequence. How and where ordinal information is represented in the primate brain remains unclear. Here, we recorded activity in populations of single neurons simultaneously from auditory cortex and ventrolateral prefrontal cortex in macaques passively listening to structured auditory sequences. Population-level analyses revealed that frontal cortex activity evolved progressively across the sequence, occupying distinct regions of neural state space for successive ordinal positions. In contrast, auditory cortex population activity reset with each element, producing stereotyped, stimulus-locked trajectories. Consistent with this dissociation, frontal cortex populations robustly decoded ordinal position independently of stimulus identity, whereas auditory cortex showed weak positional decoding despite strong identity representations. Frontal neurons further exhibited longer intrinsic timescales and reduced synchronization to element presentation, supporting integration of information across multiple elements. Together, these findings identify frontal cortex activity as a key substrate for ordinal position encoding during auditory sequence processing, while auditory cortex primarily supports element-specific sensory representations.

## Introduction

Language, music, and coordinated actions, all depend on cognitive capacity to track events. This fundamental capacity involves extracting regularities through repeated exposure and integrating information across time to track the current position within an ongoing sequence. Encoding ordinal information therefore requires representing both the identity of individual elements and their ordinal position, the latter emerging from information integrated across preceding sequence (Logan, 2021; Calmus et al., 2020). Previous studies have demonstrated potential neural mechanisms for tracking ordinal position across distributed neural networks rather than within isolated sensory regions. The frontal cortex has been repeatedly implicated in the representation of abstract sequence structure, including order (Carpenter et al., 2018; Fujii & Graybiel, 2003; Ninokura et al., 2003, 2004; Petrides, 1991; Tian et al., 2024; Xie et al., 2022), rules (Bongard & Nieder, 2010), and contextual information that generalizes across sensory features (Chen et al., 2024; Wallis et al., 2001). Recent work has shown that the frontal cortex maintains ordinal rank information in low-dimensional, rank-selective subspaces that are separable from sensory representations (Chen et al., 2024; Tian et al., 2024; Xie et al., 2022). Comparative neuroimaging work has further identified homologous frontal and temporal regions in macaques and humans engaged during auditory sequence processing (Wang et al., 2015; Wilson et al., 2015), and population recordings in macaque ventrolateral prefrontal cortex (vlPFC) during visual sequence viewing have shown that frontal populations spontaneously form joint representations of stimulus identity and abstract sequential structure even without overt task demand (Bellet et al., 2024). In contrast, the auditory cortex has been characterized primarily as encoding stimulus-specific features with high temporal precision, with single neurons in macaque auditory cortex reliably time-locking to acoustic stimuli (Kayser et al., 2010; Scott et al., 2011). While auditory cortex is sensitive to learned sequence structure and transitional probabilities (Kikuchi et al., 2017), it remains unclear whether auditory cortex maintains ordinal position representations or whether such abstract codes emerge primarily in higher-order areas.

One framework for addressing this question comes from hierarchical models, which propose that early sensory areas encode rapidly changing stimulus features while higher-order areas integrate information over longer temporal windows to represent more abstract aspects of sequence structure (Felleman & Van Essen, 1991; Kaas & Hackett, 2000). Consistent with this framework, hierarchical gradients of intrinsic timescales have been widely reported across cortex, with shorter timescales in sensory regions and longer timescales in frontal regions (Cavanagh et al., 2020; Chaudhuri et al., 2015; Gao et al., 2020; Murray et al., 2014; Wolff et al., 2022). Related but conceptually distinct gradients have also been observed in temporal integration windows within the auditory hierarchy, which increase progressively from primary auditory cortex to belt and parabelt regions and into higher-order temporal and frontal areas (Dheerendra et al., 2022; Hasson et al., 2008; Honey et al., 2012; Norman-Haignere et al., 2022; Rupp et al., 2024), and human intracranial work has revealed a cross-area gradient from early auditory regions encoding transitional probabilities to higher-order regions including the inferior frontal gyrus encoding ordinal position and identity with increasing abstraction (Henin et al., 2021). Additionally, neurons with longer timescales carry behaviourally relevant information more robustly (Runyan et al., 2017) and support more stable working memory codes in the prefrontal cortex (Cavanagh et al., 2018; Wasmuht et al., 2018), suggesting that intrinsic timescales may constraint which forms of auditory sequential information can be stably maintained in auditory hierarchy. However, evidence for whether ordinal encoding is represented in the auditory hierarchy at the population level, and how such manifestation is associated with intrinsic dynamics in the primate brain, remains limited.

Here, we simultaneously recorded populations of single neurons from auditory cortex and vlPFC in macaques during listening to learned structured of auditory sequences. We show that frontal neurons exhibit progressive population dynamics that track ordinal position, whereas auditory cortex neurons show stimulus-locked, element-specific representations. These differences are reflected in decoding performance and intrinsic properties, with frontal neurons exhibiting longer timescales and reduced synchronization to element presentation compared to auditory cortex, providing a mechanistic account for their enhanced capacity to integrate information across multiple sequence elements in order. These findings demonstrate the neural substrates of ordinal encoding through auditory sequences and show how differences in intrinsic neural dynamics constrain which forms of sequential information can be stably represented at neural populations during auditory sequence processing in the frontal and auditory cortex.

## Results

We simultaneously recorded neurophysiological data from the auditory cortex and vlPFC using linear electrode arrays acutely implanted in two adult macaques during a passive sequence learning paradigm (Figure 1; Supplementary Figure 1; methods). We isolated 676 neurons in the frontal cortex (monkey T: 371 neurons, monkey W: 305 neurons) and 798 neurons in the auditory cortex (monkey T: 413 neurons, monkey W: 385 neurons) across 79 sessions (monkey T: 31 sessions, monkey W: 48 sessions).

**Figure 1.**
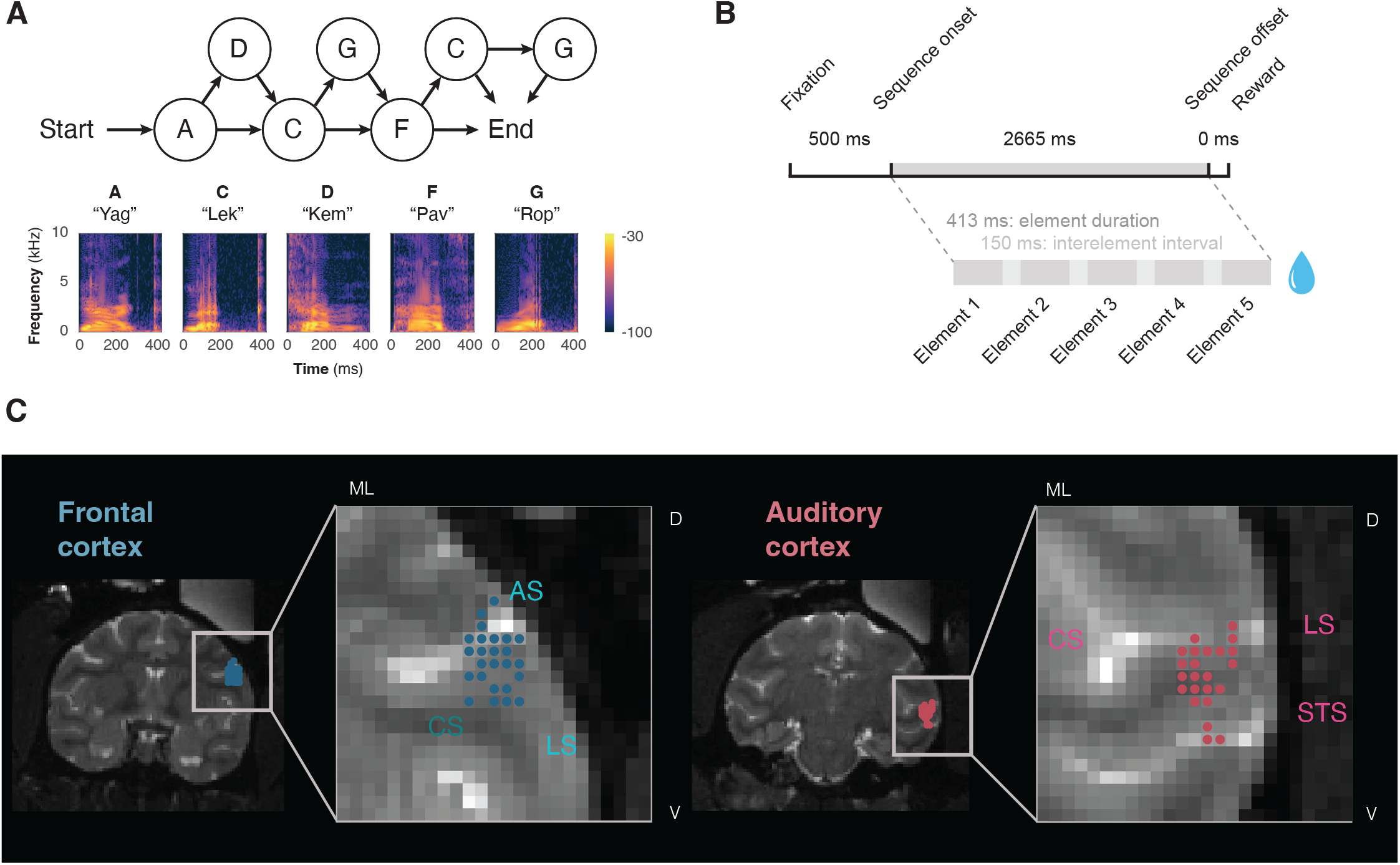
Neural recordings during a probabilistic rule-based sequence learning task. **(A)** Top: Transition graph of the artificial grammar used to generate auditory sequences. Sequences begin at node A and proceed through a series of forward-branching transitions to the end node. Bottom: Spectrograms of the five nonsense word stimuli serving as sequence elements — A (“Yag”), C (“Lek”), D (“Kem”), F (“Pav”), and G (“Rop”) — each approximately 413 ms in duration. Colour scale indicates power in dB. (B) Temporal structure of the task; after fixating for 500 ms at a central cue, a sequence was presented comprised of five acoustic elements, at the end of which reward was delivered. (C) Coronal view of recording sites in frontal (left) and auditory cortex (right) from Monkey W, presented in a representative slice. Circles represent the site corresponding to the tip of the electrode. AS, arcuate sulcus; CS, circular sulcus; LS, lateral sulcus; STS, superior temporal sulcus.

### Frontal cortex populations encode ordinal position in auditory sequences

To determine whether neural population activity preserves information about ordinal position within auditory sequences, we examined the low-dimensional dynamics of neuronal populations in frontal and auditory cortex using principal component analysis (PCA). If a population encodes ordinal position, its activity should evolve across successive elements rather than resetting with each stimulus. Consistent with this prediction, population activity in frontal cortex exhibited progressively evolving trajectories across the sequence, occupying distinct regions of neural state space for successive elements (Figure 2A, left; Supplementary Figure 2B, left). In contrast, auditory cortex population trajectories were tightly time-locked to individual elements and repeatedly returned to a similar state at each element onset (Figure 2A, right; Supplementary Figure 2B, right), indicating a reset of population activity on an element-by-element basis.

**Figure 2.**
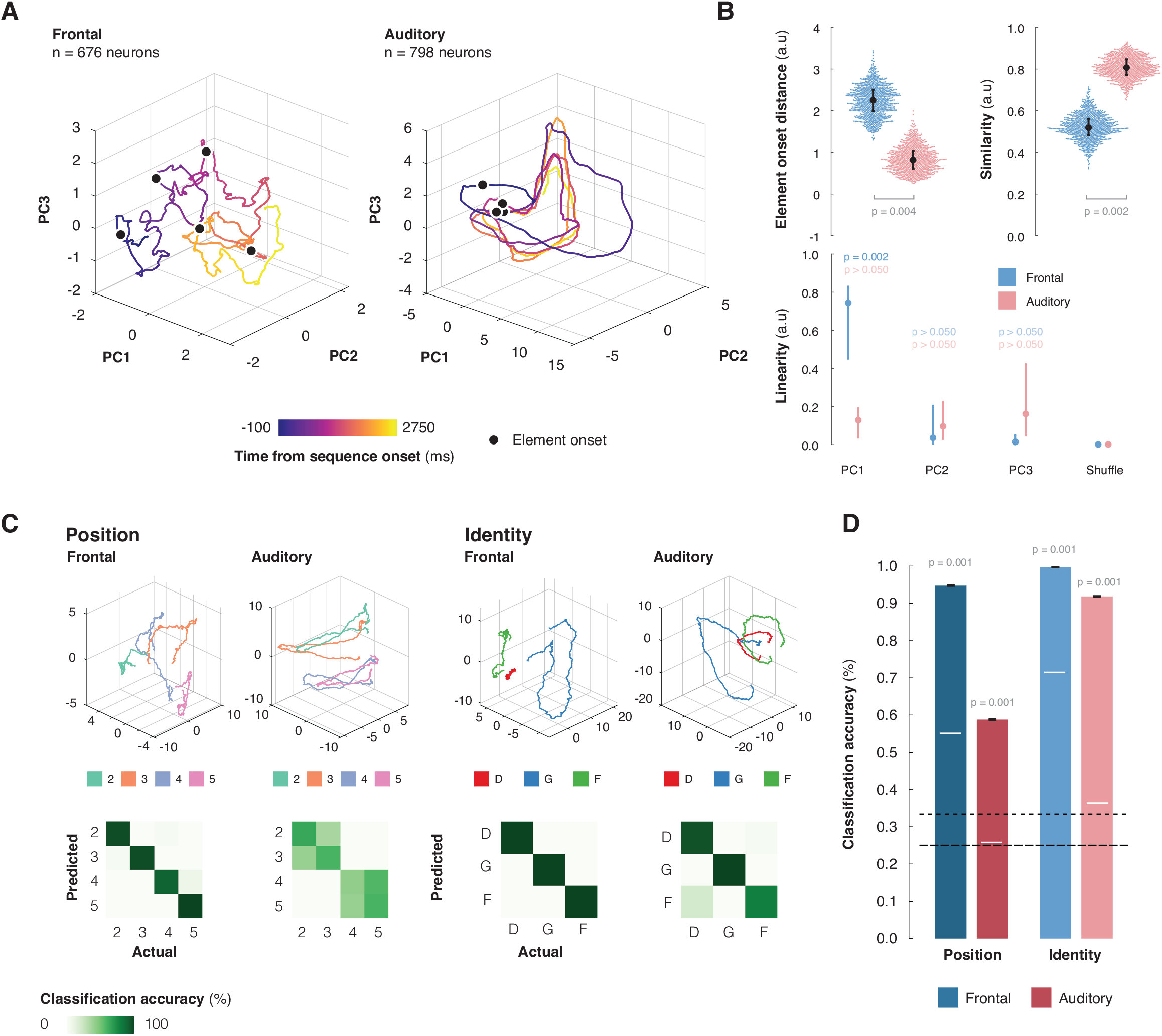
Differential neural trajectories during sequence processing in frontal and auditory cortex. (A) Principal component (PC) projections of frontal (left) and auditory (right) population activity aligned to sequence onset, showing the evolution of neural trajectories over time (add time bin size). Trajectories are color-coded by timepoint, illustrating structured temporal progression through neural state space. The onset of an element within the sequence is denoted by a black circle. (B) Differences in element-onset distance (top left), similarity (top right), and linearity (bottom) between frontal (blue) and auditory (red) population trajectories. (C) Principal component (PC) projections of frontal (sub-left) and auditory (sub-right) population activity, when averaged across position (left) or identity (right). Confusion matrices, computed from classification accuracy for each element presentation period, are shown below. (D) Bootstrapped classification accuracy for decoding element identity or position, based on PCA dynamics for auditory and frontal neural populations. Black dashed lines represent chance for position (large dash) and identity (small dash). White lines represent the classification accuracy when only using the principal component that explained the most variance.

We quantified these differences using complementary measures of population dynamics. First, if population activity resets with each element, the distance between trajectory start points should be small across elements. Auditory cortex trajectories showed significantly closer start points than frontal cortex trajectories, consistent with repeated resetting (bootstrapped pairwise difference: 1.43 [CI: 0.43 to 2.30], p = 0.004; Figure 2B, top left). Second, if population dynamics follow the same geometric pattern for each element, trajectory shapes should overlap across the sequence. Auditory cortex trajectories exhibited higher Procrustes similarity across elements than frontal cortex trajectories (bootstrapped pairwise difference: 0.288 [CI: 0.445 to 0.141], p = 0.002; Figure 2B, top right), indicating more stereotyped, element-locked dynamics. Finally, population codes that preserve ordinal position are expected to evolve monotonically with sequence progression. In frontal cortex, the dominant population trajectory showed a strong linear relationship with time in the sequence (PC1 R^2^: 0.76 [CI: 0.04 to 0.89], p = 0.002; Figure 2B, bottom; Supplementary Figure 2B), whereas no comparable relationship emerged among the three top auditory cortex components explaining the greatest variance. (all p > 0.05). This monotonic evolution indicates that frontal cortex population activity preserves information about sequence progression rather than resetting at each element.

Together, these results demonstrate that frontal cortex population carry robust information about the position of items within auditory sequences.

### Ordinal position decoding is robust in frontal cortex, but weak in auditory cortex

If frontal cortex population dynamics preserve information about sequence progression, then ordinal position should be decodable from population activity independently of stimulus identity. To test this, we used linear discriminant analysis to quantify how well neural population activity in frontal and auditory cortex, epoched around the element onset, distinguished the ordinal position of a stimulus element when identity was held constant. In frontal cortex, ordinal position was decoded with high accuracy, indicating robust representation of position within the sequence (94.84% [CI: 92.16 to 96.97%], p < 0.001; Figure 2C, D). In contrast, position decoding from auditory cortex populations was substantially weaker, although still significantly above chance (58.57% [CI: 54.54 to 62.77%], p < 0.001), indicating limited sensitivity to ordinal position when stimulus identity was controlled. Decoding accuracy for position was significantly greater in frontal cortex than auditory cortex (bootstrapped comparison: p < 0.001).

We next asked whether this difference reflected a general limitation in auditory cortex population coding or a selective difference in positional information. When position was held constant and classifiers were trained to decode stimulus identity, decoding accuracy was high in both auditory cortex (92.03% [CI: 87.91 to 94.57%], p < 0.001) and frontal cortex (99.84% [CI: 98.94 to 100.00%], p < 0.001; Figure 2C, D). Thus, both regions robustly encoded stimulus identity, whereas reliable encoding of ordinal position was selectively expressed in frontal cortex populations.

To determine how ordinal and identity information were distributed across the previously identified population dimensions, we examined decoding performance across successive principal components. In frontal cortex, ordinal position was decoded more accurately from PC2–3 than from PC1 alone (Δ = 15.7% [CI: 3.87 to 28.33%], p < 0.001; Supplementary Figure 2A,C), indicating that positional information was not dominated by a single low-dimensional axis. Instead, decoding accuracy accumulated progressively across multiple dimensions in frontal cortex but plateaued rapidly in auditory cortex (Supplementary Figure 2D), suggesting that frontal populations encode ordinal position through a distributed, high-dimensional code, whereas auditory cortex carries weaker positional information within a smaller number of low-variance dimensions.

### Single neurons exhibit mixed selectivity for identity and position

To understand how population-level ordinal representations relate to single-neuron responses, we examined the activity of individual units during auditory sequence processing. We first identified neurons significantly modulated by sequence elements by comparing firing rates following element onset to a pre-sequence baseline. Using this criterion, we identified 323 neurons in auditory cortex (monkey T: 169; monkey W: 154) and 159 neurons in frontal cortex (monkey T: 91; monkey W: 68) that showed significant modulation to at least one identity or ordinal position (Figure 3A).

**Figure 3.**
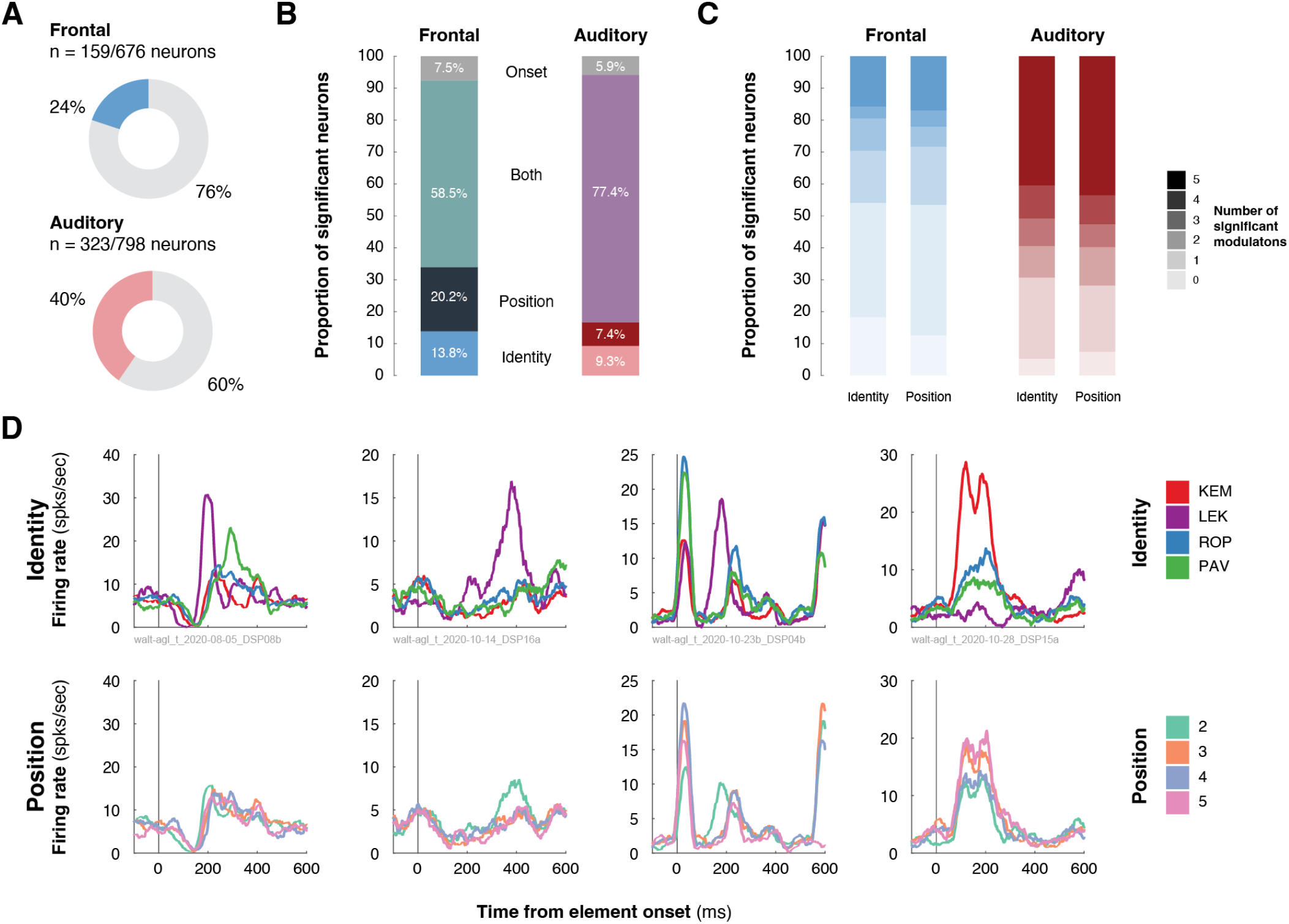
Single unit encoding of element identity and ordinal position. (A) Proportion of single neurons in auditory (top) and frontal (bottom) cortices that demonstrate significant modulation to a sequence element. (B) Proportion of significant single neurons in auditory (left) and frontal (right) cortices that demonstrate significant modulation to either: the first element in a sequence (onset, grey), a particular position in a sequence (dark), a particular element identity (light), or multiplex position and identity information. (C) Proportion of significant single neurons in auditory (left) and frontal (right) cortices that demonstrate significant modulation to a number (none to all) of elements within a sequence. (D) Example neurons aligned to element onset and averaged over element identities (top) or positions (bottom).

Across both cortical areas, single neurons commonly exhibited mixed selectivity, responding to combinations of element identity and ordinal position rather than encoding a single feature in isolation (Figure 3B, D). In auditory cortex, 77.4% of neurons (N = 250) were modulated by both identity and position, with smaller populations selective for identity alone (9.3%, N = 30), position alone (7.4%, N = 24), or sequence onset (5.9%, N = 19). In frontal cortex, 58.5% of neurons (N = 93) encoded both features, alongside distinct populations selective for identity alone (13.8%, N = 22), position alone (20.2%, N = 32), or onset responses (7.5%, N = 12). The distribution of selectivity types differed significantly between cortical areas (χ^2^ test of independence, χ^2^ (3) = 22.64, p < 0.0001), with position-only selectivity more prevalent in frontal cortex.

We next quantified the breadth of selectivity by measuring the number of distinct identities or ordinal positions to which each neuron responded (Figure 3C). Auditory cortex neurons tended to exhibit broader tuning, with many neurons responding to multiple identities and positions, including 40.6% (N = 131) responsive to all five identities and 43.7% (N = 141) responsive to all five positions. In contrast, frontal cortex neurons more frequently responded to a smaller subset of identities or positions, with significantly narrower selectivity distributions for both features (identity: Z = 6.70, p < 0.0001 position: Z = 6.43, p < 0.0001). This difference in selectivity breadth suggests that neurons in both auditory and frontal cortex carry stimulus-linked information, but that frontal cortical dynamics enable ordinal rank to emerge at the population level through more differentiated and temporally extended response patterns.

Although many neurons showed significant selectivity for sequence features, single neurons in both auditory and frontal cortex exhibited heterogeneous temporal firing patterns, including transient onset responses, sustained or ramping activity during sound presentation, and phasic offset responses (Supplementary Figure 3A). Despite this diversity, responses broadly fell into four categories: event-locked facilitated, event-locked suppressed, facilitated ramping, and suppressed ramping. Event-locked facilitated neurons exhibited transient or multiphasic increases in firing rate aligned to element onset or offset (n = 206), whereas event-locked suppressed neurons showed corresponding decreases in firing rate with similar temporal structure (n = 197). Ramping neurons displayed slow, monotonic changes in firing rate across the element duration, without clear responses to its onset or offset (Supplementary Figure 3A, n = 44 facilitated ramping; n = 35 suppressed ramping). All response classes were present in both areas, and their relative proportions were largely similar between auditory and frontal cortex (Supplementary Figure 3B; χ^2^ test of independence,χ^2^ (3) = 0.01, p = 0.9998).

### Intrinsic timescales and reduced periodicity may support ordinal integration in frontal cortex

Encoding ordinal position requires neural activity that preserves information across successive elements rather than resetting with each stimulus. We therefore asked whether differences in temporal structure and integration properties at the single-neuron level could account for the population-level ordinal encoding observed in frontal cortex.

We first examined the temporal periodicity of neuronal spiking using autocorrelation analyses. Within the sequence, elements were presented at fixed intervals (interstimulus interval: ∼563 ms; presentation rate ∼1.8 Hz). Encoding ordinal position requires neural activity that preserves information across successive elements rather than resetting with each stimulus. We therefore hypothesized that neurons supporting ordinal position encoding would show reduced periodic activity to the stimulus rhythm, reflecting temporal integration across elements. Consistent with this prediction, periodic event-locked facilitated and both types of ramping neurons in frontal cortex mostly exhibited weak and inconsistent periodic structure in their firing (Figure 4A, top; Supplementary Figure 4A). Periodicity was, however, observed in event-locked suppressed neurons, with a noticeable peak around the element presentation interval. In contrast, both event-locked facilitated and suppressed neurons in auditory cortex exhibited pronounced periodic structure, with clear positive-lag peaks aligned to the inter-element interval (Figure 4A, bottom; Supplementary Figure 4A). These findings indicate that neurons in the frontal cortex, but not auditory, do not reliably modulate in an element-by-element manner, consistent with population dynamics that evolve across the sequence.

**Figure 4.**
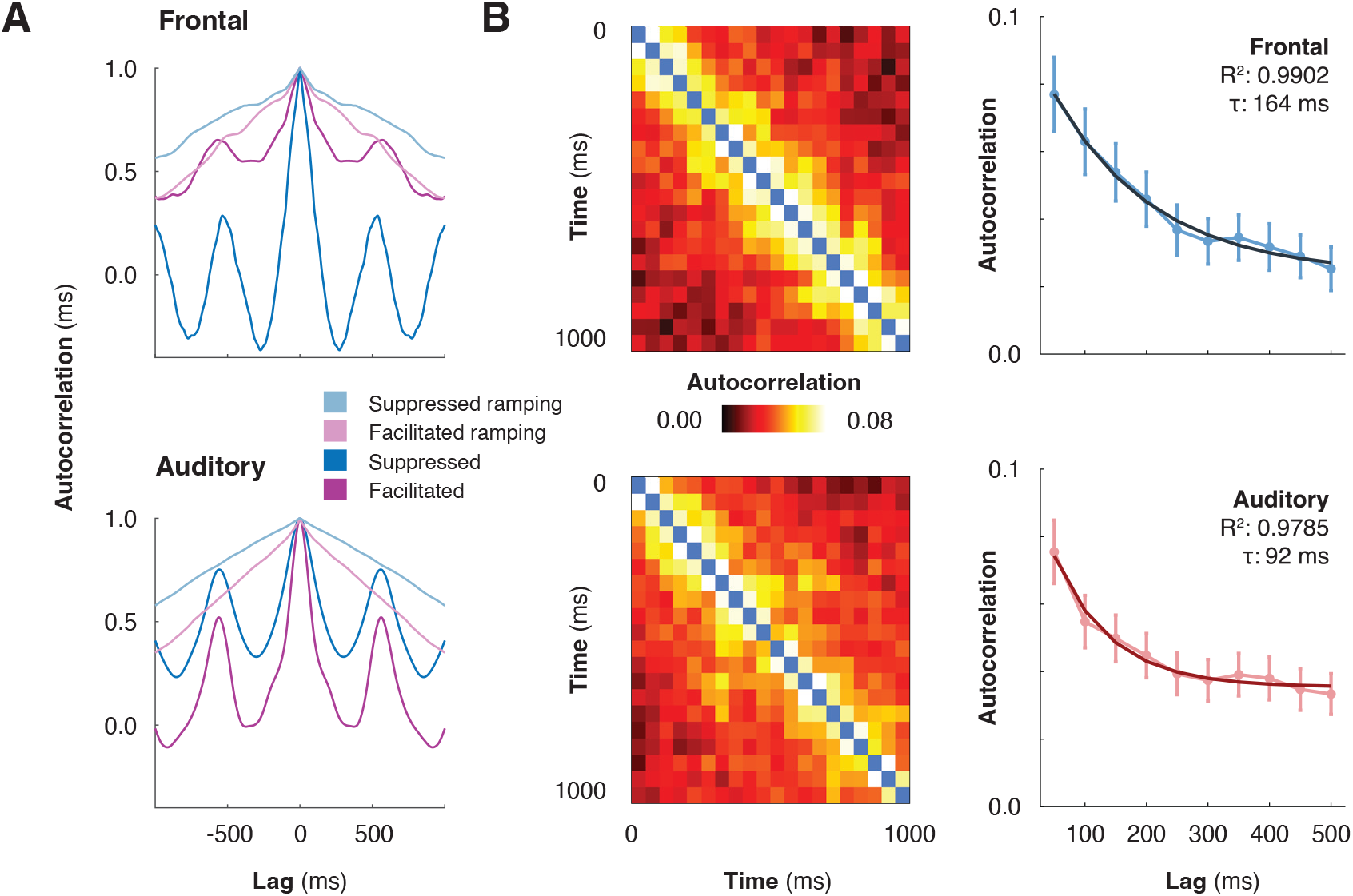
Robust periodicity in auditory cortex neurons, but not frontal cortex neurons. (A) Autocorrelation functions for auditory (left) and frontal (right) neurons, showing differences in temporal persistence of firing patterns. (B) Representative autocorrelation functions and exponential fits for auditory (top) and frontal (bottom) neurons. Insets show goodness-of-fit (R^2^) and estimated intrinsic timescale (τ).

Then, to test whether single-neuron temporal integration differs across cortical areas, we examined the intrinsic timescale of neuronal activity. We hypothesized that neurons supporting ordinal position encoding would operate on longer intrinsic timescales, enabling integration of information across successive sequence elements. Intrinsic timescale (τ) quantifies how long a neuron’s activity remains temporally autocorrelated, providing an estimate of the duration over which it integrates and retains information over time. Neurons in frontal cortex exhibited significantly longer intrinsic timescales than those in auditory cortex (frontal: 168 ms [CI: 123 to 245 ms]; auditory: 96 ms [CI: 64 to 149 ms]; difference: 73 ms [CI: 6 to 150 ms], p = 0.034; Figure 4B; Supplementary Figure 4B). By integrating information over longer temporal windows, frontal cortex neurons are well suited to preserve sequence history across multiple elements, supporting robust encoding of ordinal position during auditory sequence processing.

Together, reduced stimulus locking and longer intrinsic timescales may provide a mechanistic basis for the evolving population dynamics observed in frontal cortex. These properties enable frontal cortex neurons to integrate information across multiple sequence elements, supporting robust encoding of ordinal position during auditory sequence processing.

## Discussion

The present study provides evidence of a fundamental dissociation in how frontal and auditory cortex represent learned structured auditory sequences. Frontal population activity evolved progressively across sequences, occupying distinct regions of neural state space for successive ordinal positions, whereas auditory cortex activity reset with each element, producing stereotyped, stimulus-locked trajectories. This dissociation was reflected in decoding performance and intrinsic timescales: frontal populations robustly decoded ordinal position with longer timescales, whereas auditory cortex showed weak positional decoding despite strong identity representations. These findings indicate temporally integrative, higher-order representations of ordinal information manifested in the frontal cortex during auditory sequence processing.

The progressive evolution of frontal population activity is consistent with recent demonstrations that frontal cortex maintains ordinal rank in low-dimensional, rank-selective subspaces that are separable from sensory representations (Chen et al., 2024; Tian et al., 2024; Xie et al., 2022), and with prior work showing that frontal populations spontaneously form joint representations of stimulus identity and abstract sequential structure within distinct but superimposed neuronal subspaces (Bellet et al., 2024). Frontal population neurons representing the ordinal information were found in the vlPFC. Previous work on frontal ordinal coding have come predominantly from dorsolateral prefrontal cortex (dlPFC) using largely visual or motor sequences which require working memory components where ordinal information needed to be maintained during a delay period to guide subsequent behaviours. Our finding demonstrates that an analogous computation arises in vlPFC during task-irrelevant auditory listening when the animals were engaged in a simple visual fixation task and were never required to attend to or remember the sounds. Most prior work on frontal ordinal coding have used active tasks in which animals were trained to produce guided-behaviours by sequence orders priorly presented (Carpenter et al., 2018; Chen et al., 2024; Marc et al., 2026; Tian et al., 2024; Xie et al., 2022) in which ordinal representations could potentially reflect task-related processes such as response planning than abstract variables emerged from the sequence structure. In this study, we show that ordinal information is spontaneously constructed in frontal neuronal population activity by using sequences whose statistical structure had been internalised through prolonged exposure without any demand to encode sequence order. Our finding is aligned with and extends recent evidence that vlPFC populations represent abstract structural properties of auditory sequences without overt task demand (Bellet et al., 2024).

This is consistent with the broader role of these areas in the primate frontotemporal sequence-processing network. In this study the recordings were made from areas 44 and 45, which received projections from auditory and multisensory association cortex via the ventral temporo-frontal system (Frey et al., 2014; Gerbella et al., 2010; Romanski et al., 1999). These areas have been implicated in auditory sequencing operations that require working-memory-like maintenance of relating elements separated in time, including the processing of non-adjacent dependencies in structured auditory sequences (Calmus et al., 2020; Wilson et al., 2017). The ordinal neuronal representations we observed in areas 44/45 provide a candidate neural substrate for this kind of maintenance, complementing the working-memory-based ordinal representations described in dlPFC during active tasks (Carpenter et al., 2018; Chen et al., 2024; Tian et al., 2024). More broadly, these findings position macaque areas 44/45 as a candidate substrate for the evolutionary precursors of sequencing computations under passive listening which is closer to how naturalistic auditory sequences such as speech, conspecific vocalizations and music are encoded and processed in everyday listening, through long-term exposure rather than encoding ordinal information based on trial-by-trial basis task specific rules. In contrast, the stimulus-locked, resetting dynamics of auditory cortex suggest a fundamentally different computational role. Auditory cortex population activity produced stereotyped trajectories that reset with each element, consistent with a sensory representation that prioritizes stimulus identity over ordinal position. This is consistent with prior evidence that auditory cortex exhibits precise temporal coding and stimulus-specific responses, and with hierarchical models in which stimulus-locked dynamics in early auditory areas are transformed into more abstract representations in higher-order regions (Camalier et al., 2019; Romanski & Averbeck, 2009; Scott et al., 2011; Yin et al., 2020) (Supplementary Discussion).

The mechanistic basis for this dissociation may lie in the intrinsic temporal properties of neurons in each region. As ordinal position must be computed by integrating information across multiple successive elements, stable positional representations are expected to emerge preferentially in circuits whose intrinsic dynamics preserve information over extended temporal intervals (Calmus et al., 2020; Logan, 2021). Hierarchical gradients of intrinsic timescales have been widely reported, with shorter timescales in sensory cortices and longer timescales in frontal regions (Cavanagh et al., 2020; Chaudhuri et al., 2015; Gao et al., 2020; Murray et al., 2014; Wolff et al., 2022). In this study, we demonstrated that auditory cortex neurons exhibited pronounced periodic firing at the inter-element interval, reflecting rhythmic resetting of neural activity. Frontal cortex neurons, by contrast, showed little evidence of such periodicity. Instead, they displayed significantly longer intrinsic timescales and gradual ramping dynamics that scaled with sequence progression rather than locking to individual element onsets, consistent with a neural architecture that preserves a running representation of the sequence across elements from which positional information can be derived (Supplementary Discussion).

In summary, we showed a functional dissociation of ordinal representations between auditory and frontal cortex during processing of learned auditory sequences, consistent with hierarchical accounts of cortical organization in which early sensory areas encode rapidly changing stimulus features while higher-order association cortices integrate information over longer temporal windows (Hasson et al., 2008; Murray et al., 2014; Norman-Haignere et al., 2022), and with predictive coding models in which higher-order regions form predictions about internalized complex structures of sequences (Dehaene et al., 2015; Heilbron & Chait, 2018). Beyond the cortical regions examined here, the hippocampus is a compelling candidate for this integrative role across longer timescales given its contribution to statistical learning and relational structure (Henin et al., 2021; Zhou & Turk-Browne, 2025), Interactions among sensory, frontal and hippocampal systems may contribute to the neural foundation for higher-order cognitive functions, including language processing, music perception, and complex action planning.

## Materials and Method

### Animals and ethical statement

Two adult male rhesus macaques (Macaca mulatta; Monkey T: 12.3 kg, 11 years old; Monkey W: 9.4 kg, 8 years old) from a group-housed colony were studied. All macaque procedures performed in this study were approved by the UK Home Office (project license: 70/8318) and by the Animal Welfare and Ethical Review Body at Newcastle University. The work complies with the Animal Scientific Procedures Act (1986) on the care and use of animals in research and with the European Directive on the protection of animals used in research (2010/63/EU). We support the principles of the consortium on Animal Research Reporting of In Vivo Experiments (ARRIVE).

### Behavioural paradigm

The artificial grammar learning (AGL) paradigm was used in this study. The task was originally designed to examine probabilistic auditory sequence structure acquired through implicit learning (Attaheri et al., 2015; Kikuchi et al., 2017; Wilson et al., 2013). Briefly, nonsense word sounds were arranged into sequences that were consistent with the allowed AG transitions between words or violated these transitions based on the artificial grammar of Saffran et al. (1996). The transition graph (Figure 1A, top) consists of multiple forward-branching relationships and has considerable variability in the transitional probabilities. The paradigm consists of five nonsense words serving as sequence elements (A, C, D, F, and G): “yag,” “lek,” “kem,” “pav,” and “rop” (Figure 1A, bottom). The nonsense words were produced by a female human talker and digitally recorded using an Edirol R-09HR sound recorder (Roland Corp.) in a sound-attenuating acoustical chamber. The mean duration of the nonsense words was 413.6 ms (yag: 413.7 ms; kem: 413.1 ms; lek: 414.0 ms; pav: 413.7 ms; rop: 413.5 ms). The power spectral density of the nonsense word stimuli fell well within the macaque audible range, being at least 30 dB above hearing threshold in the range of 100-5000 Hz (Pfingst et al., 1975). All nonsense words were normalized to the maximum root-mean-square (RMS) amplitude across the stimulus set. Sequences were matched in duration (2665 ms) and contained a fixed number of elements (5 words).

Each session consisted of a fixed 30-min exposure phase in which the animals passively listened to 160 repetitions of the exposure sequences. During the exposure phase, a set of exposure sequences that followed the ordering relationships were presented in correct order following the AG. This exposure phase was followed by a testing phase (160 trials, ∼30 minutes) where both correct and violation sequences were used. During this phase, a randomly chosen sequence was individually presented over the audio speakers while the animals fixated on a spot on the monitor (all the sequences used in this study are shown in Supplementary Figure 1). Ordinal position was independent of the violation manipulation and therefore dissociable from local transitional probabilities.

The animals had previously been trained to perform a fixation task during sound stimulation, which was assessed using an infrared eye tracker (Arrington Research, AZ, US). All recordings were performed after stable sequence statistics had been established, allowing us to examine neural representations in the absence of ongoing learning. Following this, they initiated an experimental trial by fixating on a spot at the centre of the monitor for 500 ms, triggering the presentation of a randomly selected testing sequence from the sixteen testing sequences available. The macaque was required to fixate on the spot for 3,000 ms (within a ∼2–4°-diameter fixation acceptance window centred on the fixation spot) until the end of the testing sound sequence presentation to obtain a juice reward. They received the juice reward at the offset of the fixation spot for correctly fixating throughout the entire sequence presentation. Reward delivery occurred following a jittered interval after the final element in the sequence presentation, and varied between monkeys and sessions (Monkey T: 400-700 ms, uniform distribution; Monkey W: 0 to 1500 ms, normal distribution)

### Stimuli presentation

Sound waveforms were amplified, attenuated, and delivered through audio speakers at ∼75 dB SPL. The sound level was calibrated at the animal’s head position using an XL2 sound level meter (NTI Audio) before every recording session. The electrophysiological recording sessions were conducted in a darkened, foam-lined, single-walled acoustical chamber (IAC). The animals had previously been acclimated to periods of head immobilization using operant training with positive reinforcement. The macaque was seated in a recording chair with the head immobilized, facing a computer monitor approximately 60 cm in front of the subject (24” Samsung, LCD). Two free-field speakers (Creative Inspire T10) were horizontally positioned at approximately 30° visual angle. Stimulus presentation, behavioural control and reward administration were controlled with an in-house program written in Python 2.7, which is partly based on PsychoPy (Peirce, 2007) on Linux Ubuntu 16.04 LTS platform (Intel Core i3 processor; Toshiba, 4GB RAM) via a DAQ-LabJack U6-Pro Interface. Auditory stimuli were presented via the Advanced Linux Sound Architecture (ALSA).

### Neurophysiology

MRI was used for guiding chamber placement and the electrodes. The MRI structural and functional data were obtained with a 4.7T scanner (Bruker BioSpin, Etlingen, Germany). A customized, MRI-compatible head post and oval recording chamber (44.0 mm x 31 mm, PEEK) were implanted under aseptic conditions during general anaesthesia. The recording chamber was positioned stereotaxically over the right hemispheres of both animals to target the auditory and frontal cortex based on the MRI.

Multiple 16-channel electrode arrays (V-probe, Plexon, Dallas, TX) were used for the extracellular electrophysiological recordings. Stainless steel guide tubes (23 ga or 26 ga) were used to penetrate the dura matter to protect the electrode tips and to prevent deflection. After the guide tubes were in place, the electrodes were independently advanced using a remote-controlled, multichannel microdrive system (NAN-SYS-4; Plexon. Inc., Dallas, TX). The signal was referenced to the guide tube or electrode shaft (V-probe), amplified, high-pass filtered (NeuraLynx: 600-8000 Hz; TDT: 300-5000 Hz), digitized (NeuraLynx: 32000 Hz; TDT: 24414.0625 Hz), and recorded via a TDT RZ5 system for Monkey W and Digital Lynx SX for Monkey T (Neuralynx, Cheetah 5.6 software) acquisition systems.

Neural recordings obtained from vlPFC were located on the inferior convexity, encompassing areas 44 and 45 (Figure 1C). Recordings in auditory cortex were performed along the lateral sulcus, primarily targeting regions corresponding to the auditory core and belt areas (rostral temporal area [R], anterior lateral belt [AL]) on the supratemporal plane. Anatomical landmarks (lateral sulcus) and search stimuli (including tones, noise, and complex sounds) were used to ensure that the auditory cortex was reached. However, the recorded locations, neuronal spiking activity, and LFPs were not selected by their stimulus response preference nor the shape of the waveforms of the spiking activity. Throughout the recording sessions, spiking activity was aurally monitored through headphones (HD 280 Professional, Sennheiser). Single neurons were extracted offline, identified using automated spike sorting via Kilosort 4, and manually curated in Phy thereafter (Pachitariu et al., 2024). Manual curation involved evaluating cluster quality metrics (spike count, signal-to-noise ratio, and inter-spike interval violations) together with visual inspection of waveform shape and stability to ensure well-isolated single units (neurons) were retained.

### Data analysis

All data analyses were performed using custom scripts implemented in MATLAB (MathWorks, Natick, MA), although we leveraged established toolboxes including FieldTrip for some aspects of the neural data preprocessing and analysis (Oostenveld et al., 2011), and the gramm toolbox for data visualization (Morel, 2018). Only correct, rewarded, non-violation trials were included in all analyses unless otherwise stated.

All statistical comparisons between bootstrap distributions were performed using a two-sided bootstrap test. For a given pair of distributions, a difference distribution was computed (observed minus reference), and a p-value was calculated as twice the minimum of the proportion of differences greater than or equal to zero and the proportion less than or equal to zero. Confidence intervals on the difference were reported as the 2.5th, 50th, and 97.5th percentiles of the difference distribution. Bootstrap analyses used 1000 resampling iterations unless otherwise stated. For comparisons of encoding category proportions and functional cluster distributions, chi-square tests of independence were applied to contingency tables. For comparisons of ordinal count data, Wilcoxon rank-sum tests were used. A significance threshold of α = 0.05 was applied throughout.

### Decoding identity and positional coding in auditory and frontal populations

Neuronal spike density functions (SDFs) were computed by convolving spike trains with a post-synaptic potential kernel defined by:

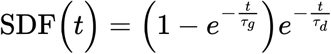

with growth time constant (τg) of 1 ms, and decay time constant (τd) of 20 ms, corresponding to the values measured for excitatory post-synaptic potentials. Firing rates were smoothed and averaged across trials, aligned to the onset of each sequence element. Baseline firing rates were calculated from a pre-stimulus window spanning 150 to 50 ms before sequence onset by averaging activity across all valid trials within that interval. SDFs were z-scored relative to this baseline to normalise activity.

Smoothed, z-scored SDFs were obtained using a moving average filter (window size = 100 ms). SDFs were computed separately for five auditory stimulus identities (A, C, D, F, G) at each of the five ordinal positions within the sequence. To evaluate positional coding independently of stimulus identity, SDFs for stimulus identity ‘C’ were extracted across four positions (2nd, 3rd, 4th, and 5th), as this was the only element that could appear in each position (Supplementary Figure 1). SDFs for identity decoding were extracted for identities C, F, and G at a fixed ordinal position (position 5). This design allowed dissociation of neural representations of stimulus identity and serial position. An analysis window of 0–413 ms post-element onset was used for all decoding analyses.

Dimensionality reduction was performed using principal component analysis (PCA) on the condition-averaged neural activity matrices (neurons × time). Condition-specific SDFs were first concatenated across conditions to form a joint matrix, and PCA was applied to this pooled representation so that all conditions shared a common low-dimensional subspace. Only neurons with valid responses in all conditions were retained, ensuring a consistent neuron set across the subspace.

Cumulative variance explained by the first six principal components was inspected for each area. The first three principal components were projected onto a three-dimensional state space, and trajectories were visualised as time-coloured curves, with element onset times marked. The first three principal components were retained as features for all decoding analyses. Linear discriminant analysis (LDA) classifiers were implemented to predict stimulus identity or ordinal position from these PC features. Classifier performance was initially assessed using a holdout cross-validation procedure (70% training, 30% testing). To robustly estimate decoding accuracy and confidence intervals, 1000 bootstrap resamples were generated with replacement. For each resample, LDA classifiers were trained on the resampled training set (70% of samples, drawn with replacement) and evaluated on the held-out unique samples. Performance metrics included classification accuracy and confusion matrices, normalised by true class counts. Statistical significance was assessed by comparing bootstrap accuracy distributions against chance-level performance (1/3 for identity, 1/4 for position) using two-tailed bootstrap tests. Confidence intervals for accuracy estimates were defined as the 2.5th to 97.5th percentiles of the bootstrap distribution.

Several geometric metrics were computed on bootstrapped PCA trajectories to characterize how population activity evolves over time. The following metrics were extracted from each bootstrapped sample: element onset difference, similarity, and linearity.

The Euclidean distance between consecutive element onsets in PC state space was computed as a measure of how far population activity had evolved between successive elements. The mean 3D PC coordinate was computed within a ±10 ms window centred on each element onset, and the inter-element onset distance was defined as the Euclidean norm of the difference between the mean coordinates of consecutive element pairs (elements 1–2, 2–3, 3–4, 4–5). The mean distance across the four consecutive pairs was taken as the summary metric for each bootstrap iteration.

Trajectory shape similarity across sequential elements was assessed using Procrustes analysis. For each bootstrap iteration, the 3D PC trajectory segment corresponding to each sequence element was extracted, and Procrustes distances between consecutive element segments were computed with scaling disabled. A Procrustes similarity index was defined as 1/(1 + d), where d is the Procrustes distance, such that values approaching 1 indicate high shape similarity.

Linearity of temporal progression in state space was assessed by computing the coefficient of determination (R^2^) of a linear regression of each of the first three PC scores against time. This analysis was performed under bootstrap resampling (1000 iterations, n = 500 neurons sampled with replacement) to obtain a distribution of R^2^ values for auditory and frontal populations separately. An identical analysis was applied to a temporally shuffled control condition in which the time axis of each neuron’s SDF was randomly permuted prior to PCA, providing a within-neuron null distribution that preserved firing rate distributions while destroying temporal structure. Statistical comparisons between observed and shuffled R^2^ distributions, and between auditory and frontal populations, were performed using a two-sided bootstrap test.

### Determining single-unit element-related responses to identity and position

We implemented a generalized linear modeling (GLM) approach to determine whether single-unit firing rates were significantly modulated by stimulus identity (i.e., sound) or presentation order. The analysis was performed on spike density functions (SDFs) time-locked to stimulus onset; for each trial, we extracted a continuous SDF and normalized it by computing z-scores based on a 100 to 0 ms baseline window relative to sound onset. Baseline normalization was performed within each sound condition to account for sound-specific variability in pre-stimulus firing. We then computed a time-resolved representation of the z-scored SDFs using a sliding window approach: 100 ms windows were averaged and shifted in 10 ms steps, yielding a trial-by-time matrix of firing rates across smoothed windows. This representation was used for time-resolved GLM fitting.

At each time window, a linear regression model was fitted to predict firing rate from the following categorical predictors: stimulus identity (A, C, D, F, G), ordinal position within the sequence (positions 1–5). Trial number was included as a dummy-variable. The regression model took the form:

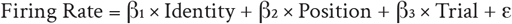

At each time point, we performed an ANOVA (using the anova function in matlab) to determine significant main effects of sound and order. To assess whether a unit encoded sound identity or order information, we examined the resulting time series of p-values for each factor within a post-stimulus analysis window (0 to 413 ms). A neuron was considered significantly modulated by a factor if there was a contiguous period of at least 50 ms (i.e., 5 or more adjacent time bins, each 10 ms apart) during which the p-value was below 0.05. This duration-based threshold ensured that only sustained effects were considered significant and reduced the likelihood of detecting transient spurious fluctuations in firing rate. The binary significance flags were recorded separately for sound identity and order position.

Neurons were classified as identity-encoding, position-encoding, both, or onset-only based on whether the GLM encoding flags were positive for any of the identity-related or position-related model terms, respectively. Neurons were assigned to the ‘onset-only’ category if they were significantly modulated overall but showed no significant identity or position encoding across the analysis window. The proportion of neurons in each encoding category was quantified separately for auditory and frontal cortex. A chi-square test of independence was applied to a 2 × 4 contingency table (area × encoding category) to assess whether the distribution of encoding types differed between cortical regions. The number of sounds significantly encoded per neuron was also compared between areas using a Wilcoxon rank-sum test.

### Clustering common patterns of single unit modulation

To identify distinct temporal response profiles among sound-responsive neurons, we clustered neurons based on the shape of their trial-averaged, baseline-normalized responses to sound stimuli. Neuronal responses were represented as z-scored SDFs aligned to sound onset, spanning 200 ms to +800 ms. These SDFs were lightly smoothed (10 ms kernel) to reduce noise while preserving temporal resolution. For each neuron, responses were averaged across all sound conditions to yield a single vector capturing its canonical response time course.

Pairwise dissimilarity between neurons was quantified using correlation distance (1 r), where r is the Pearson correlation between z-scored response profiles. The resulting symmetric distance matrix was submitted to classical multidimensional scaling (MDS; cmdscale) to obtain a low-dimensional embedding of neuronal response profiles in a 5-dimensional space. The quality of this embedding was assessed using a Shepard plot of original versus embedded distances, and non-metric MDS stress was computed as a supplementary measure (mdscale, stress criterion). The number of clusters was determined empirically by applying k-means clustering (k = 2 to 20, with 20 replicates per k) to the MDS coordinates and computing the mean silhouette coefficient for each k; the value of k maximising the mean silhouette was selected. Final cluster assignments were obtained by running k-means with 50 replicates at the chosen k. The quality of each neuron’s assignment was assessed by computing R^2^ between the neuron’s SDF and its cluster-average response profile. Clusters were grouped posthoc into four functional categories based on the shape of cluster-average responses: facilitated (transient excitation), suppressed (transient inhibition), facilitated ramping (sustained or increasing excitation), and suppressed ramping (sustained or increasing inhibition). The proportion of auditory and frontal cortex neurons assigned to each functional category was compared using a chi-square test of independence.

### Determining periodicity in auditory and frontal cortex neurons

To assess the temporal structure of neural responses during auditory sequence processing (Figure 4), we performed autocorrelation analyses on spike density functions (SDFs) aligned to sequence onset. For each neuron, SDFs were smoothed using a 100 ms moving average. Baseline-normalized SDFs were calculated by z-scoring relative to the pre-stimulus period (–200 to 0 ms). Autocorrelation was computed over a post-stimulus window spanning 0 to 2665 ms (five 563 ms auditory elements), using MATLAB’s “xcorr” function with coefficient normalization.

To identify significant recurrent structure, we extracted local maxima (peaks) in the population-averaged autocorrelation curves using MAT-LAB’s “findpeaks” function. We restricted detection to positive lags, focusing on forward-in-time recurrence, and applied a minimum peak prominence of 0.25. The first significant peak beyond zero lag was used as an estimate of the dominant recurrence time within each cluster, reflecting the timescale of temporal patterning in neural responses.

### Estimating intrinsic timescales in auditory and frontal cortex neurons

To estimate intrinsic neural timescales, we analysed trial-by-trial spike count correlations across consecutive time bins during a pre-stimulus baseline period ( −1000 to 0 ms relative to sequence onset). Spike times were binned in 50 ms intervals, yielding 20 bins per trial. This analysis was limited to neurons deemed to be significantly responsive to the element. For each significant neuron, spike counts were aggregated across trials within each bin, and Pearson correlation coefficients were computed between all pairs of bins, yielding a 20 × 20 correlation matrix reflecting the similarity of firing rate fluctuations across time lags. Bins with zero mean firing across trials were excluded from analysis to avoid spurious correlations.

The autocorrelation function (ACF) was derived by averaging correlations along diagonals of the matrix corresponding to increasing time lags (0 to 950 ms in 50 ms steps). Average ACFs were computed separately for auditory and frontal cortex neuron populations. To quantify the intrinsic timescale, the average Awith growth time constanCF was fit over lags of 50–500 ms with an exponential decay function of the form:

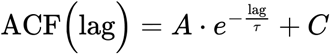

where A is the amplitude, τ is the intrinsic timescale (decay constant), and C is an offset. Nonlinear least-squares fitting was performed using MATLAB’s “lsqcurvefit” function with initial parameter estimates optimised for convergence. The fitted decay constant τ provides an estimate of the characteristic timescale over which neural activity fluctuations remain correlated, reflecting the temporal integration properties of each population. Goodness-of-fit was assessed by computing R^2^ between the empirical and fitted ACF.

## Acknowledgements

This work was supported by MRC (MR/X003701/1), BBSRC (BB/ Z00005X/1), and Brain & Behavior Young Investigator Award (33751). Animal colony maintenance was supported by Wellcome Trust (WT092606AIA, PhD studentship for Ryan Calmus) and the European Research Council (MECHIDENT). We thank Fabien Balezeau for MRI support, and Jennifer Nacef and the technical staff at the Comparative Biology Centre at Newcastle University for their support with animal husbandry, Ryan Calmus for funding acquisition, project administration or resources, Colline Poirier, Benjamin Slater, Valentina Giuffrida, Emma Woolgar, Kazutaka Takahashi and Yaneri Ayala for discussions or comments on the manuscript.

## Author contributions

Conceptualization: S.E., Y.K.; Funding: Y.K., C.I.P., T.D.G., S.E.; Data curation: S.E., J.B., Y.K.; Formal analysis: S.E.; Investigation: B.A., Y.K.; Methodology: S.E., Y.K; Supervision: Y.K., C.I.P., T.G.; Visualization, S.E.; Writing - original draft, S.E., Y.K.; Writing - review and editing: S.E., B.A., J.B., T.G., C.I.P., Y.K.

