## Supplementary Info for "Ordinal position in auditory sequences is encoded in population dynamics in macaque frontal cortex"

### SUPPLEMENTARY FIGURES

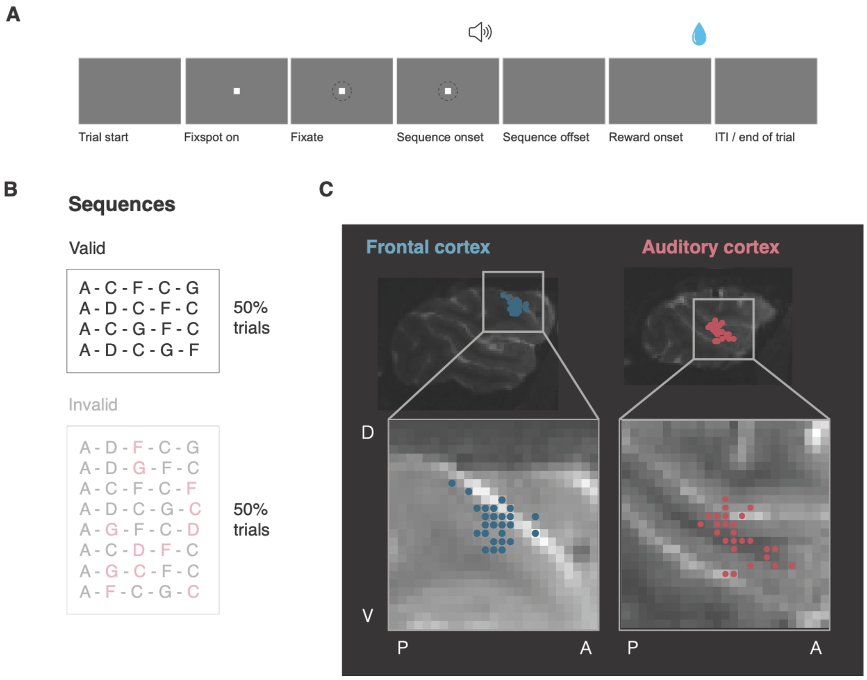


**Supplementary Figure 1 | Auditory sequence structure and transitional probabilities**. **(A)** Example task structure. **(B)** Structural schematic of the auditory sequences used in the study. The sequences, with their order following the rule indicated by arrows (Figure 1 A), are shown in the top panel; the sequences that violated this structure (bottom panel) were not analysed in this manuscript. The highlighted elements in the invalid sequences denote where the first violation occurs. **(C)** Sagittal view of recording sites in frontal (left) and auditory cortex (right) from Monkey W, presented in a representative slice. Circles represent the site corresponding to the tip of the electrode.

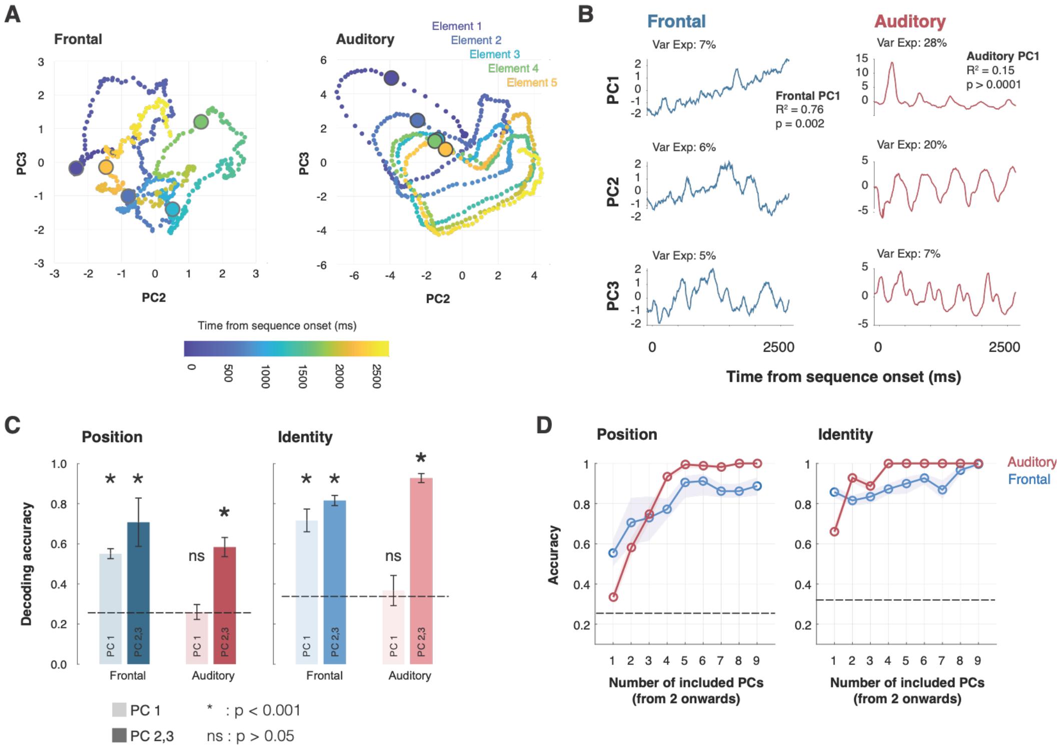


**Supplementary Figure 2 | Principal components of population activity**. **(A)** PC2 and PC3 of neuronal population activity for the frontal (left) and auditory (right) cortex, plotted as a function of time from sequence onset (shown through point colour). **(B)** The first three principal components (PC1–PC3) of neuronal population activity for frontal (left, blue) and auditory (right, red) cortex, plotted as a function of time from sequence onset. Variance explained is noted above each PC. **(C)** Bootstrapped classification accuracy for decoding element identity or position, based on PCA scores for either PC1 alone (light) or PC2&3 (dark) combined, for both auditory (red) and frontal (blue) neural populations. **(D)** Bootstrapped classification accuracy for decoding element identity or position, based on the number of included PCs (excluding PC1) in the linear discriminant analysis, for frontal (blue) and auditory (red) populations. Frontal populations encode position through a distributed, high-dimensional code, whereas auditory cortex carries weaker positional information across fewer low-variance dimensions.

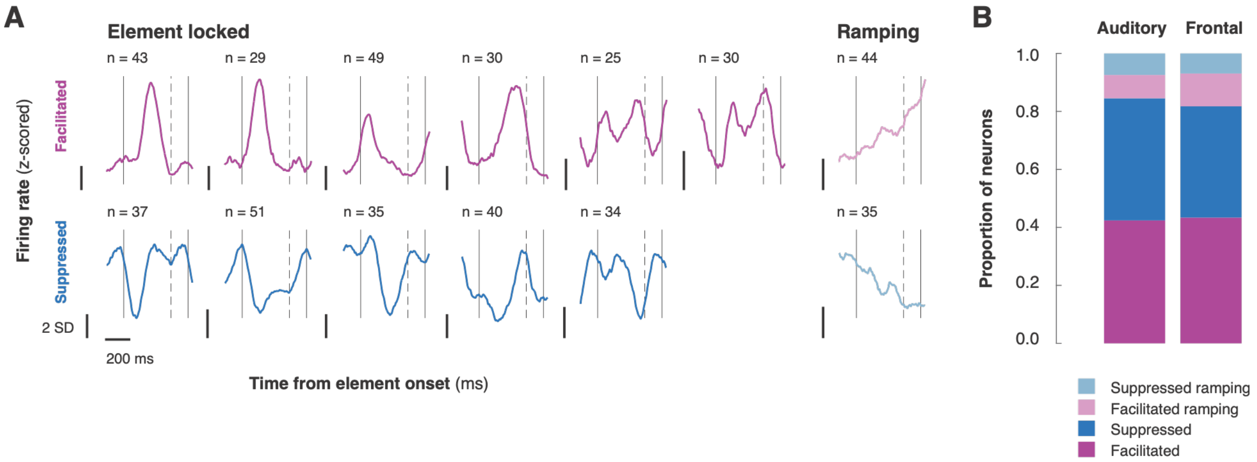


**Supplementary Figure 3 | Diversity of single-neuron temporal response profiles**. **(A)** Average element-aligned spike density functions of different single-unit response classes identified across both regions. These patterns were qualitatively grouped into facilitated, suppressed, facilitated ramping, and suppressed ramping (**B**) Proportion of neurons belonging to each response class in the auditory and frontal cortex. The relative distribution of facilitated, suppressed, and ramping dynamics was largely similar across both cortical areas.

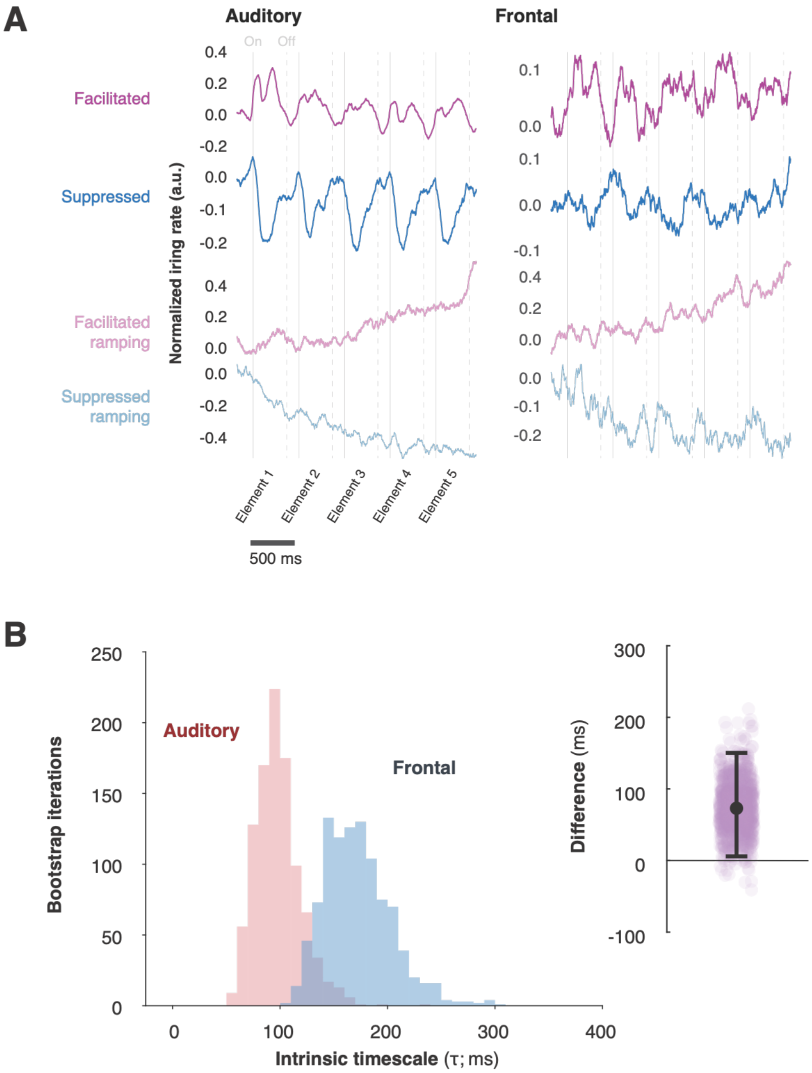


**Supplementary Figure 4 | Differences in intrinsic timescales between cortical areas**. **(A)** Average sequence spike density functions of different single-unit response classes identified across both regions. (**B**) Bootstrapped distributions of the intrinsic timescale for auditory (red) and frontal (blue) cortex. *Inset*: mean difference in intrinsic timescale between the two regions, with the 95% confidence interval.

### SUPPLEMENTARY DISCUSSION

#### Ordinal and identity decoding from neuronal populations in auditory and frontal cortex

Both auditory and vlPFC carried strong identity information in the full population response, but the underlying coding structure differed. In auditory cortex, identity information was concentrated in a small number of non-dominant components, which decoded identity more accurately than the corresponding components in frontal cortex (**Supplementary Figure 2C, D**). In frontal cortex, by contrast, identity information was distributed across many dimensions, including the dominant component (PC1), aligning with prior evidence on decoding accuracy within the auditory ventral stream (Russ et al., 2008). We note that our auditory recordings were from core and belt areas; identity selectivity in higher auditory areas may be even more concentrated and selective (Fukushima et al., 2014; Kikuchi et al., 2010). Although full-population decoding revealed weaker ordinal position coding in auditory cortex, principal component analysis showed that ordinal information is not absent but subordinate to the dominant sensory response (**Supplementary Figure 2B**). When the dominant variance component was excluded, ordinal position became decodable from the remaining dimensions (**Supplementary Figure 2C**). The structure of this positional code differed fundamentally between regions. Frontal cortex distributed ordinal information across many population dimensions, whereas auditory cortex concentrated it in a small number of low variance components (**Supplementary Figure 2D**). This dimensionality difference was mirrored at the single-neuron level, where we observed that position-selective neurons were significantly more prevalent in frontal cortex than in auditory cortex (**Figure 3B**). The greater prevalence and diversity of position-selective responses in frontal cortex is consistent with the mixed selectivity characteristic of prefrontal populations, which generates the high-dimensional population codes capable of flexibly representing abstract task variables.

This gradient — from subordinate, low-dimensional positional signals in sensory cortex to dominant, high-dimensional positional representations in frontal cortex —aligns with human intracranial recordings, with early brain areas tracking lower-level sensory inputs and transitional probabilities, while the inferior frontal gyrus, anterior temporal lobe and hippocampus encode higher-order aspects of sequences such as ordinal position and identity (Henin et al., 2021). Our findings also align with the observation that content (frequency) and ordinal position (structure) are encoded and maintained as dissociable representations in the human auditory system, possessing distinct characteristics and can be reactivated independently (Fan et al., 2021). Auditory cortex nonetheless is not entirely insensitive to sequence structure: learned ordering relationships modulate auditory cortical oscillatory coupling and transient responses under passive exposure of auditory sequences (Kikuchi et al., 2017), but such sensitivity likely reflects encoding of local transitional probability changes rather than global ordinal position tracking.

#### Interpreting ramping activity in the frontal cortex

A potential alternative interpretation is that the progressive frontal dynamics reflect reward expectation associated with the fixation task rather than ordinal position. Several features of our data argue against this. Frontal population activity did not evolve smoothly across the trial but occupied discrete, element-aligned regions of state space for successive ordinal positions, and ordinal position was decodable from activity time-locked to individual sound onsets (**Figure 2C, D**) – neither of which would be expected from a monotonic reward anticipation signal. Ordinal decoding generalized across multiple well-learned sequences whose elements differed at matched ordinal positions, indicating a code for position within sequence rather than for the identity of specific elements. Auditory cortex was recorded simultaneously, in the same animals and under identical fixation and reward conditions yet showed no comparable positional code in the full population response.

Principal component analysis provided further evidence against the interpretation that ordinal coding is reducible to reward anticipation. In frontal cortex, the dominant variance component (PC1) exhibited a strong linear trajectory across ordinal positions (**Figure 2B, Supplementary Figure 2B**), consistent with a signal that covaries with elapsed sequence time, which may include reward-anticipation components known to be present in prefrontal cortex (Stoll & Rudebeck, 2024). Critically, however, ordinal position was decoded more accurately from frontal PC2-3 than from PC1 alone (**Supplementary Figure 2C**), and successive positions occupied distinct, non-collinear regions in the PC2-3 subspace (**Supplementary Figure 2A**). This demonstrates that frontal populations carry ordinal position information in dimensions independent of any scalar signal associated with elapsed time or anticipation of reward delivery, and that the multidimensional population code described here cannot be reduced to a single anticipatory ramp.
